# Short-chain carboxylates facilitate the counting of yeasts in Sub-high temperature *Daqu*

**DOI:** 10.1101/2023.12.22.573164

**Authors:** Zhiqiang Ren, Juan Xie, Tuoxian Tang, Zhiguo Huang

**Author notes:** Corresponding author, Zhiqiang Ren;, Zhiguo Huang;.

## Abstract

Sub-high temperature Daqu is a traditional solid fermenting agent that produces Chinese strong-aroma Baijiu. It is abundant in various microorganisms, including bacteria, yeasts, molds, and actinomycetes, of which yeasts play a crucial role in ethanol production and flavor formation. Counting yeasts in Daqu is difficult due to the interference of molds and bacteria. Antibiotics are employed to inhibit bacterial growth, but there is no effective way to suppress molds without affecting the growth of yeasts. In this study, short-chain carboxylates (C1-C6) were added to the culture medium at various pH conditions to investigate their effects on the growth of molds and yeasts. Results showed that they have distinct inhibitory effects in a pH-and concentration-dependent manner. A few of these tested short-chain carboxylates effectively suppress mold growth in agar plates without affecting yeast growth. Herein, a simple and feasible method for improving the efficiency of yeast isolation and counting in Daqu has been proposed, which may be useful for studying yeasts from complex habitats.

## 1. Introduction

Sub-high temperature Daqu (STD) is a saccharifying and fermenting agent made from wheat, which is often used to produce Chinese strong-aroma Baijiu. It plays a key role in fermentation [1–2], including the degradation of complex natural substrates such as starch and protein, and the transformation of degraded products into alcohols, acids, esters, aldehydes, and other flavor substances 3, which dictates the quality and specialty of Baijiu brewing 14. STD is produced by natural inoculation, which is abundant in microorganisms, including a large number of molds, yeasts, and bacteria, as well as a small number of actinomycetes [5–6]. In a recent study that investigated the community structure of fungi in STD using second-generation sequencing technology, it was shown that yeasts accounted for 60% of the total fungi 7. A total of 420 fungal strains were isolated from 30 Daqu starter samples using the plate culture method and identified using ITS region sequencing. Of these isolated strains, 386 (92%) were yeasts, and 34 (8%) were filamentous fungi. *Saccharomyces cerevisiae*, *Wickerhamomyces anomalus*, and *Saccharomyces fibuligera* were the dominant species, accounting for 79% of the relative abundance 8. Yeasts are the primary microorganisms that convert sugars into ethanol during the fermentation process and participate in the production of some volatile flavor compounds 9. Therefore, many studies focus on the whole yeast population and isolated yeast colonies in STD [10–12].

It is tricky to effectively isolate and count yeasts in STD due to the interference of other microorganisms, especially molds 13. Supplementing antibiotics in culture media can effectively inhibit bacterial growth and circumvent the interference of bacteria 14. Using the classical plate culture method, sixteen pure yeast cultures and various yeast strains with special functions were successfully isolated from STD samples [15–17]. However, the number of pure yeast species obtained through classical methods is significantly lower than those identified by high-throughput sequencing. To isolate and analyze the whole population of yeasts, it is necessary to find a way to inhibit the growth of molds and keep yeast growth uncompromised.

Previous studies have shown that organic acids inhibit the growth of molds and yeasts. At concentrations between 0.5 and 2.5 g/L, valerate, propionate, and butyrate completely suppress mold growth, while a higher concentration of acetate and lactate is needed to achieve similar inhibitory effects [18–20]. Many organic acids were shown to inhibit yeast growth, such as formic acid 21, acetic acid 22, propionate, lactate 19, and caproic acid 23. Likewise, these organic acids inhibit yeast growth in a concentration-dependent manner. It has been reported that yeasts can grow naturally when the lactate concentration is below 100 mM, but when the lactate concentration reaches 400 mM, the growth rate of yeast decreases by 50% 23.

Even though it is well-established that adding organic acids to the culture medium regulates the growth of molds and yeasts, few studies have been conducted to investigate whether organic acids show different effects on the growth of molds and yeasts at different pH values and concentrations. This study aims to investigate the effects of short-chain carboxylates (C1-C6) on the growth of yeasts and molds in sub-high temperature Daqu with anticipation of finding a culture condition that favors the growth of yeasts and inhibits the growth of molds, allowing efficient isolation and counting of yeasts from a mixed culture. As far as we know, this is the first study that investigates the inhibitory effects of various short-chain carboxylates on molds and yeasts in the sub-high temperature Daqu.

## 2. Materials and Methods

### 2.1 Sample collection

Sampling was carried out in the strong-flavor Baijiu production factory in Yibin, Sichuan Province, China. Daqu for production was collected from ten fermentation workshops and 0.5 kg samples were collected from each workshop on November 3, 2022. The samples collected were a mixture of small particles and powder. After mixing and milling, samples were passed through a 60-mesh screen and then transferred to sterile bags, sealed and frozen at −20 °C, and shipped to the Sichuan University of Science and Engineering, Yibin, China on dry ice for further use.

### 2.2 Yeast culture media

Malt extract agar (MEA) 24 was prepared by dissolving 20 g malt extract, 10 g glucose, 5 g peptone, and 20 g agar in 1 L distilled water. For potato dextrose agar (PDA) 25, 15 g of potato extract, 20 g glucose, and 20 g agar were dissolved in 1 L distilled water. For rose Bengal agar (RBA) 26, 5 g peptone, 10 g glucose, 1 g potassium dihydrogen phosphate, 0.5 g magnesium sulfate (MgSO_4_·7H_2_O), 100 ml of a 1/3,000 aqueous solution of rose Bengal, 20 g agar were dissolved in 1 L distilled water. For Wallerstein laboratory nutrient agar (WL) 27, 4 g yeast extract, 5 g tryptone, 50 g glucose, 0.425 g potassium chloride, 0.125 g calcium chloride, 0.125 g magnesium sulfate (MgSO_4_·7H_2_O), 0.55 g potassium dihydrogen phosphate, 0.0025 g ferric chloride, 0.0025 g manganese sulfate (MnSO_4_·H_2_O), 0.022 g bromocresol green, 20 g agar were dissolved in 1 L distilled water. For yeast extract peptone dextrose agar (YPD) 28, 10 g yeast extract, 20 g peptone, 20 g glucose, and 20 g agar were dissolved in 1 L distilled water. When the sterilized culture medium was cooled to approximately 50 °C, antibiotics chloramphenicol was added to all culture medium to the final concentration of 50 μg/mL. Short-chain carboxylates were added to the various media at the desired concentrations, and the pH value of the medium was adjusted by the addition of 1 M HCl or 1 M NaOH, as required by the test. Diluent saline peptone (SPO) was prepared by dissolving 8.5 g sodium chloride, 0.3 g disodium hydrogen phosphate (Na_2_HPO_4_·12H_2_O), and 1 g peptone in 1 L distilled water. SPO was adjusted to pH 5.6 by the addition of 1 M HCl and 1 M NaOH.

### 2.3 Enumeration

Samples (10 g) were mixed with 90 mL SPO, soaked at 4 °C for 30 min, and homogenized with a Vortex Genie2 (Scientific Industries, America) at ‘10’ speed (2 min), duplicate counting plates were prepared using appropriate dilutions. For spread-plating, 0.1 mL of the dilution was spread on the surface of a dried plate. After incubation, the colonies appearing on the plates were counted and calculated as colony-forming units (CFU) per gram Daqu sample. For plates covered with molds that cannot be counted directly by the naked eye, the plates are inverted on a strong light source, a photograph is taken, and the counting is performed.

### 2.4 Data analysis

Yeast counting numbers from three independent experiments are presented as means ± SEM. Statistical significance was determined by a two-sided unpaired t-test. p<0.05 was the significance threshold. All statistical analysis was performed using GraphPad Prism 8.0 (GraphPad Prism Software Inc., La Jolla).

## 3. Results and discussion

### 3.1 MEA is used as a yeast culture medium

Microbial suspensions of STD with three dilutions were plated onto five types of commonly used yeast culture media, including MEA, PDA, RBA, WL, and YPD. When the dilution factor is 100, a large number of yeast colonies grow on plates, which makes yeast counting difficult. What’s more, too much mold growth interferes with the yeast counting (Figure 1A). When the dilution factor is 1,000, a few hundred yeast colonies can be counted, and the interference of molds is attenuated (Figure 1A). More importantly, the number of yeast colonies in this range met the criteria of yeast isolation and counting, which ranged from 30 to 300 CFU 29. Only a few yeast colonies can be counted when the microbial suspension is diluted 10,000 times, which is far below the counting and culture criteria (Figure 1A). Therefore, a dilution factor of 1,000 was used in subsequent experiments.

**Figure 1.**
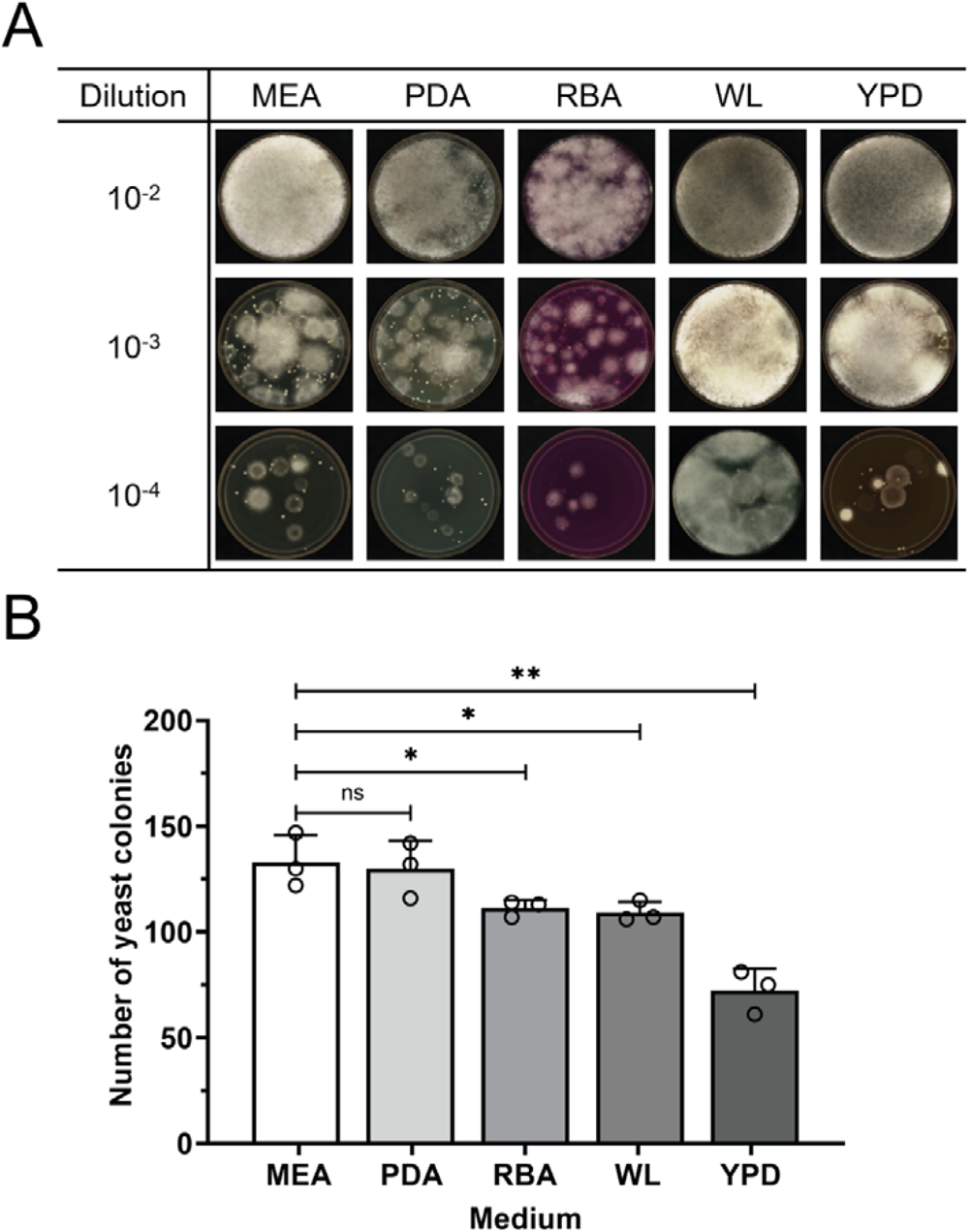
The fungal growth of STD in different yeast culture media. A, representative agar plates; B, yeast colony counting numbers at 1,000 times dilution of Daqu. MEA, malt extract agar; PDA, potato dextrose agar; RBA, rose Bengal agar; WL, Wallerstein laboratory nutrient agar; YPD, yeast extract peptone dextrose agar. 10^-2^: 100 times dilution; 10^-3^: 1,000 times dilution; 10^-4^: 10,000 times dilution. ns: no statistically significant difference; *: statistically significant difference, p<0.05; **: statistically significant difference, p<0.01.

Of the five tested yeast culture media, MEA showed the highest number of yeast colonies, followed by PDA, RBA, WL, and YPD (Figure 1B). There was no statistically significant difference in the number of yeast colonies between MEA and PDA. The main nutrients in MEA come from malt juice which is a good match with the raw materials used to prepare STD. Accordingly, MEA was selected as the growth medium for isolating and counting yeasts.

### 3.2 The effects of formate and propionate on the growth of yeasts and molds

In this study, the MEA medium was supplemented with short-chain carboxylates at concentrations of 0.05 M, 0.1 M, and 0.2 M, and the pH of the medium was adjusted in appropriate ranges to study their effects on the growth of molds and yeasts. For the medium that is supplemented with formate, the pH was adjusted in a range of 4.2 to 5.2. Both the pH value and formate concentration showed effects on the growth of molds and yeasts. When the pH is lower than 4.8, little biomass can be found on agar plates (Figure 2A). Molds and yeasts started to grow at pH 4.8 and more biomass was found at pH 5.0 and 5.2. Agar plates with a higher pH value showed a larger number of yeasts. When the final concentration of formate added in the medium was 0.05 M, the number of yeast colonies at pH 4.8, 5.0, and 5.2 was (2.17 ± 0.51) × 10^5^ CFU/g Daqu, (8.43 ± 0.21) × 10^5^ CFU/g Daqu, and (12.17 ± 0.21) × 10^5^ CFU/g Daqu, respectively (Figure 2C). Furthermore, formate inhibited the growth of molds and yeasts in a concentration-dependent manner. When the agar plates were at pH 5.2 and the concentrations of formate in the medium were 0.05 M, 0.1 M, and 0.2 M, the number of yeast colonies was (12.17 ± 0.21) × 10^5^ CFU/g Daqu, (8.37 ± 0.87) × 10^5^ CFU/g Daqu, and (2.30 ± 0.26) × 10^5^ CFU/g Daqu, respectively (Figure 2C).

**Figure 2.**
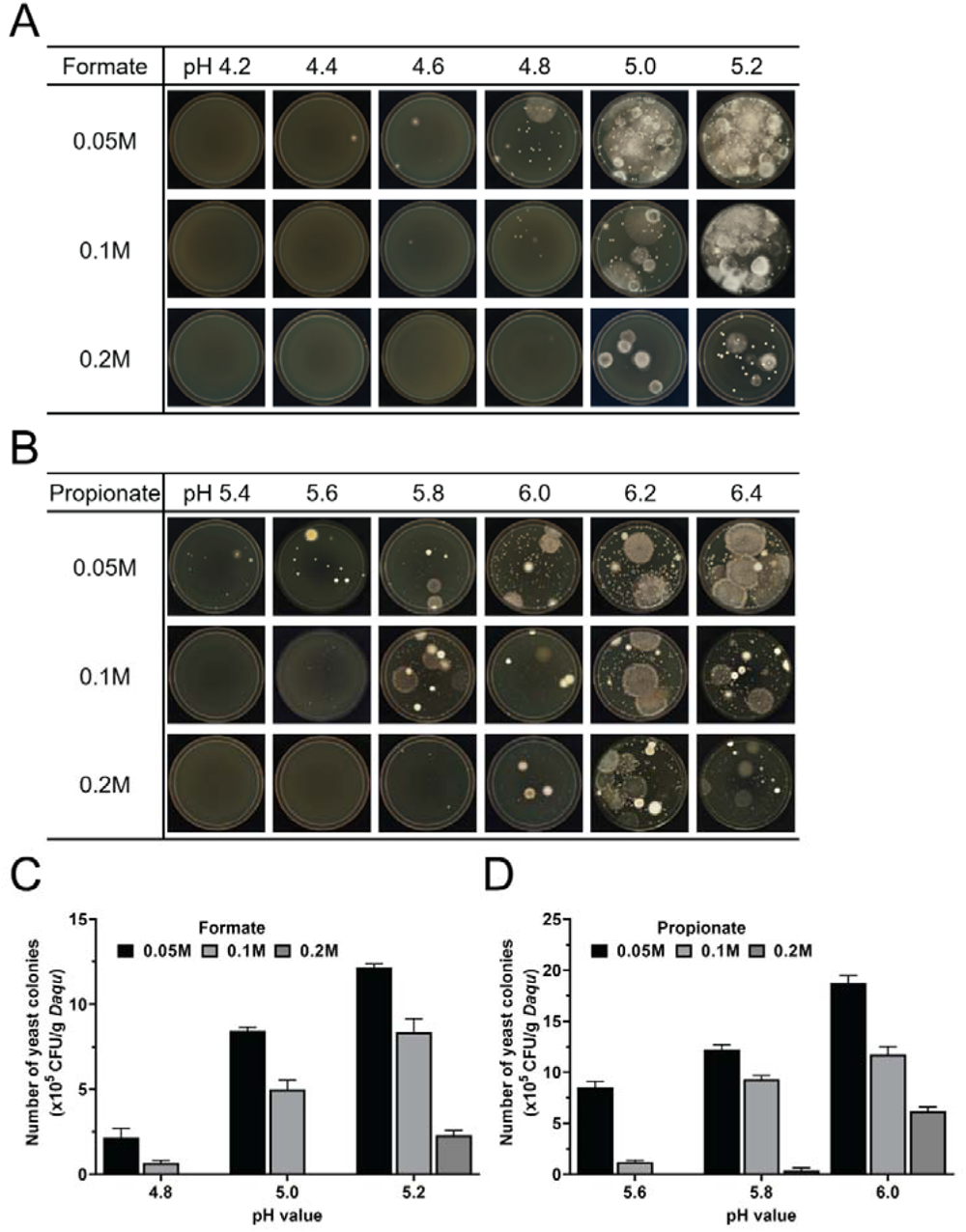
The effect of formate or propionate on the fungal growth of STD. A, representative agar plates at different pHs and supplemented with different amounts of formate; B, representative agar plates at different pHs and supplemented with different amounts of propionate; C, yeast colony counting numbers (CFU) from the agar plates at different pHs and supplemented with different amounts of formate; D, yeast colony counting numbers (CFU) from the agar plates at different pHs and supplemented with different amounts of propionate.

For the medium that is supplemented with propionate, the pH was adjusted in a range of 5.4 to 6.4. Molds and yeasts started to grow at pH 5.4 and more biomass was found at pH 6.0, 6.2, and 6.4 (Figure 2B). When the final concentration of propionate added in the medium was 0.05 M, the number of yeast colonies at pH 5.6, 5.8, and 6.0 was (8.53 ± 0.55) × 10^5^ CFU/g Daqu, (12.23 ± 0.47) × 10^5^ CFU/g Daqu, and (18.73 ± 0.75) × 10^5^ CFU/g Daqu, respectively (Figure 2D). Consistent with the observations from the agar plates that were supplemented with formate, higher concentrations of propionate resulted in higher cellular toxicity to the yeasts. When the agar plates were at pH 6.0 and the concentrations of propionate in the medium were 0.05 M, 0.1 M, and 0.2 M, the number of yeast colonies was (18.73 ± 0.75) × 10^5^ CFU/g Daqu, (11.77 ± 0.76) × 10^5^ CFU/g Daqu, and (6.17 ± 0.40) × 10^5^ CFU/g Daqu, respectively (Figure 2D).

### 3.3 The effects of acetate and butyrate on the growth of yeasts and molds

It has been reported that acetate and butyrate affect the growth of yeast and mold 20[30–31]. In this study, for the medium that is supplemented with acetate, the pH was adjusted to a range of 4.8 to 5.8. Molds predominate in the microflora that grew on the agar plates when the pH is higher than 5.4 (Figure 3A), while lower pH values favor the growth of yeast colonies and strongly the growth of mold. Only yeast colonies were observed when the pH value of agar plates reached 4.8 and 5.0 (Figure 3A). A high concentration of acetate suppressed the growth of yeasts. When the agar plates were at pH 5.0 and the concentrations of acetate in the medium were 0.05 M, 0.1 M, and 0.2 M, the number of yeast colonies was (15.70 ± 0.20) × 10^5^ CFU/g Daqu (10.90 ± 0.85) × 10^5^ CFU/g Daqu, and (7.50 ± 0.44) × 10^5^ CFU/g Daqu, respectively (Figure 3C).

**Figure 3.**
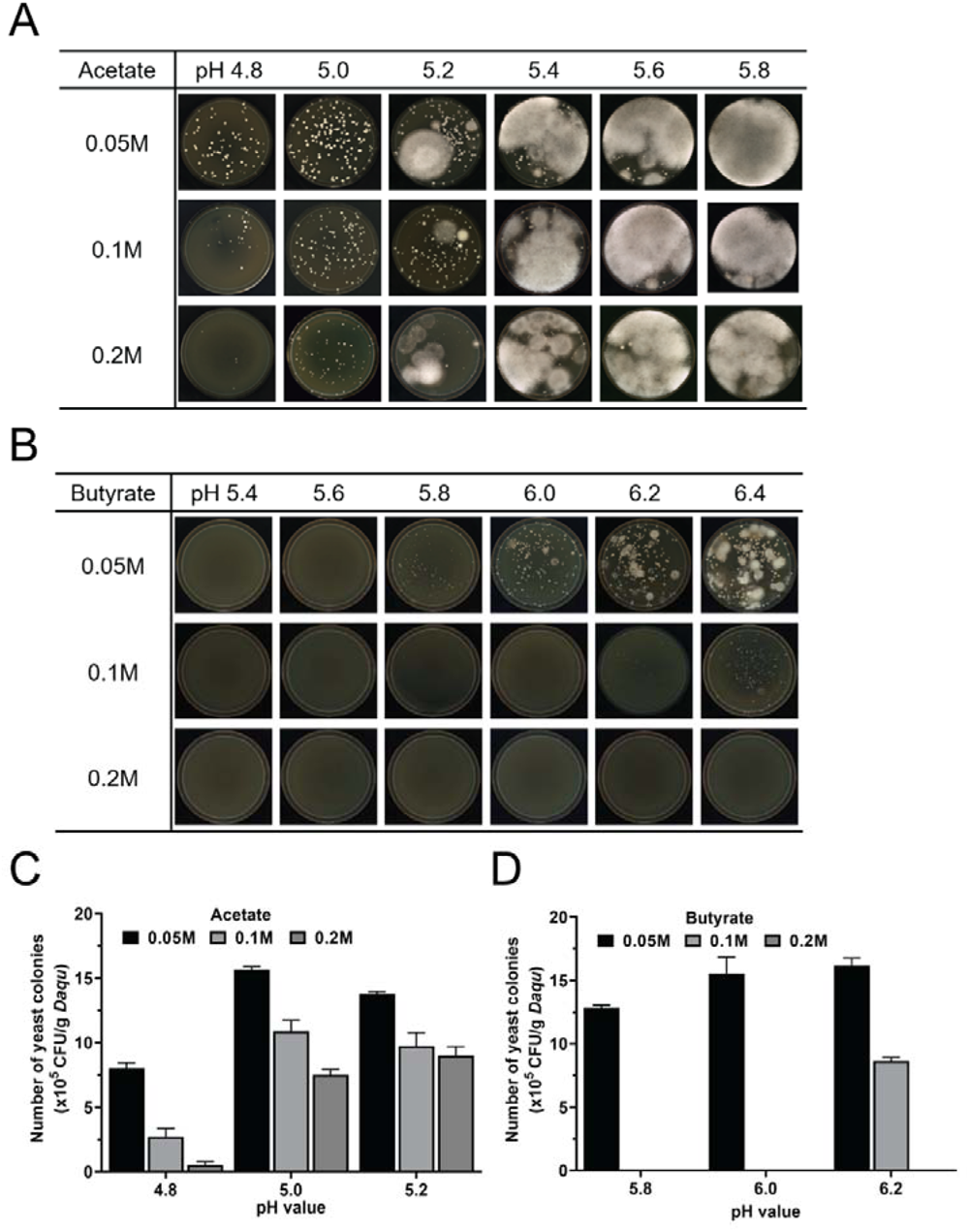
The effect of acetate or butyrate on the fungal growth of STD. A, representative agar plates at different pHs and supplemented with different amounts of acetate; B, representative agar plates at different pHs and supplemented with different amounts of butyrate; C, yeast colony counting numbers (CFU) from the agar plates at different pHs and supplemented with different amounts of acetate; D, yeast colony counting numbers (CFU) from the agar plates at different pHs and supplemented with different amounts of butyrate.

Butyrate showed stronger cellular toxicity compared with acetate. In a pH range of 5.4 to 6.4, no yeast colonies or mold were observed when the butyrate was added at a concentration of 0.2 M (Figure 3B). Yeasts and molds started to grow when the pH value of the medium was 5.8 and the final concentration of butyrate in the medium was 0.05 M. For the agar plates that were supplemented with butyrate, higher pH values favor microbial growth. When the final concentration of butyrate added in the medium was 0.05 M, the number of yeast colonies at pH 5.8, 6.0, and 6.2 was (12.83 ± 0.21) × 10^5^ CFU/g Daqu, (15.53 ± 1.32) × 10^5^ CFU/g Daqu, and (16.17 ± 0.61) × 10^5^ CFU/g Daqu, respectively (Figure 3D).

Acetate and butyrate exhibited different effects on microbial growth in STD. At pH 5.8, a large number of molds grew on the agar plates supplemented with acetate, while only yeast colonies were observed on the agar plates supplemented with butyrate, which indicated that besides their distinct cellular toxicity, short-chain carboxylates may regulate the growth of yeasts and molds through altering metabolic pathways. The growth pattern of yeasts and molds could be modulated so that molds were suppressed and yeasts were unaffected, which facilitates the study of yeasts in mixed culture.

### 3.4 The effects of lactate and pyruvate on the growth of yeasts and molds

Lactate and pyruvate are common organic acids produced during microbial fermentation. Previous studies showed that they have some effects on the growth of molds and yeasts [32–33]. For the medium that is supplemented with lactate, the pH was adjusted to a range of 3.0 to 4.4. Molds appeared on all agar plates, which indicated that lactate can’t effectively inhibit mold growth (Figure 4A). At low pH values, a higher concentration of lactate inhibited the growth of yeasts. When the agar plates were at pH 3.0 and the concentrations of lactate in the medium were 0.05 M, 0.1 M, and 0.2 M, the number of yeast colonies is (2.67 ± 0.42) × 10^5^ CFU/g Daqu, (0.60 ± 0.10) × 10^5^ CFU/g Daqu, and (0.03 ± 0.06) × 10^5^ CFU/g Daqu, respectively. However, a higher concentration of lactate did not have stronger inhibitory effects on yeast growth when the agar plates were at pH 3.4 (Figure 4C).

**Figure 4.**
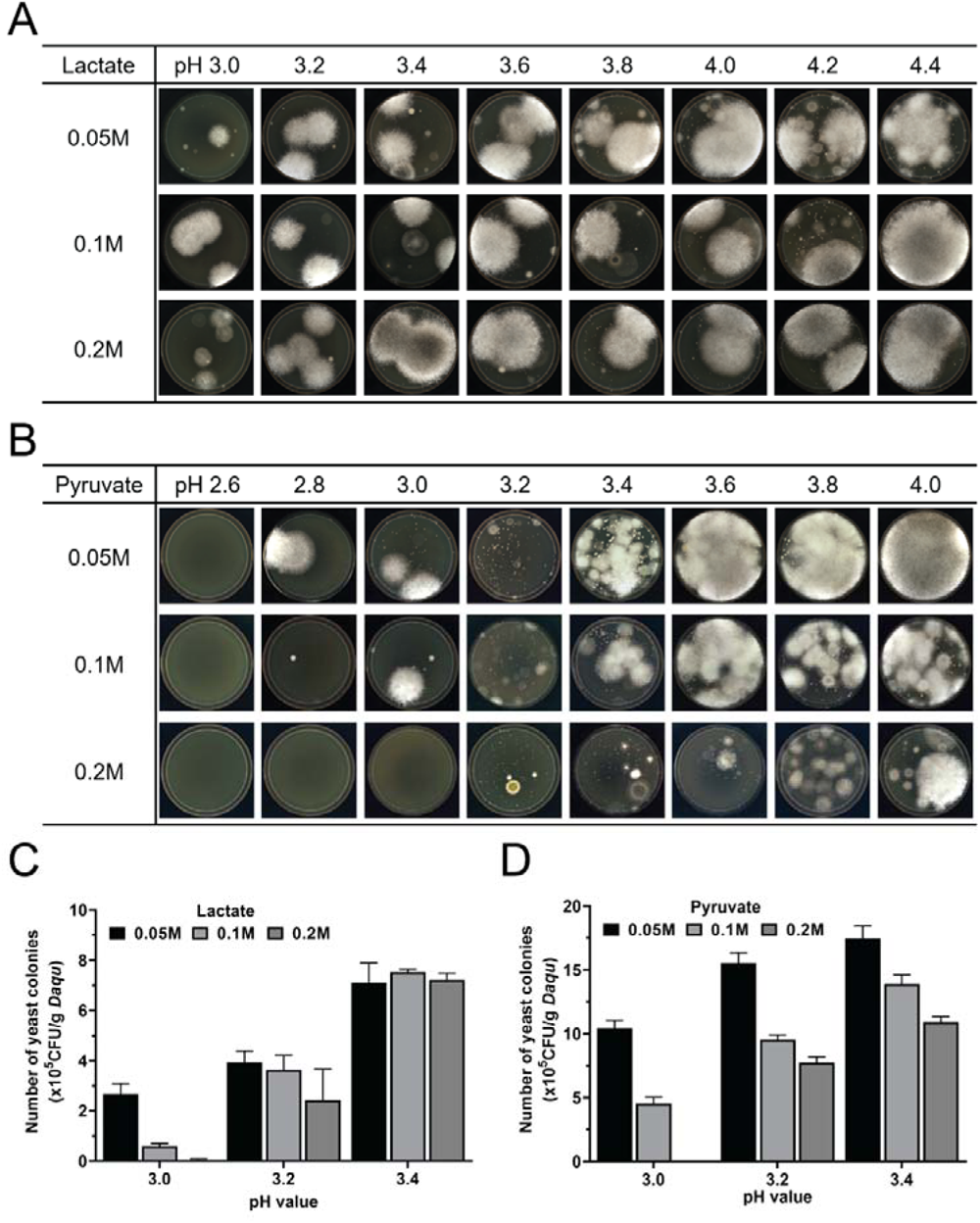
The effect of lactate or pyruvate on the fungal growth of STD. A, representative agar plates at different pHs and supplemented with different amounts of lactate; B, representative agar plates at different pHs and supplemented with different amounts of pyruvate; C, yeast colony counting numbers (CFU) from the agar plates at different pHs and supplemented with different amounts of lactate; D, yeast colony counting numbers (CFU) from the agar plates at different pHs and supplemented with different amounts of pyruvate.

For the medium that is supplemented with pyruvate, the pH was adjusted to a range of 2.6 to 4.0. At pH 2.6 and 2.8, little biomass was found on agar plates. The number of yeast colonies started to increase at pH 3.0, but molds predominate in the microbial growth at higher pH values (Figure 4B). Consistent with what has been found in other short-chain carboxylates, pyruvate inhibited the growth of yeast in a concentration-dependent manner. As the concentration of pyruvate increases, the number of yeast colonies decreases. When the agar plates were at pH 3.2 and the concentrations of pyruvate in the medium were 0.05 M, 0.1 M, and 0.2 M, the number of yeast colonies was (15.53 ± 0.81) × 10^5^ CFU/g Daqu, (9.53 ± 0.38) × 10^5^ CFU/g Daqu, and (7.73 ± 0.47) × 10^5^ CFU/g Daqu, respectively (Figure 4D). Interestingly, molds and yeasts grew at a very low pH when the agar plates were supplemented with lactate or pyruvate, in contrast to the agar plates that were supplemented with other short-chain carboxylates, where no molds or yeasts grew when the pH was below 4.2.

### 3.5 Short-chain carboxylates have different inhibitory effects on fungal growth

To compare the inhibitory effects of short-chain carboxylates on the growth of yeasts and molds in STD, the MEA medium was supplemented with formate, acetate, propionate, butyrate, valerate, caproate, lactate, and pyruvate in a final concentration of 0.05 M, and the pH of media was adjusted to 5.7 to match the natural pH of MEA. A large amount of molds grew on the agar plates supplemented with formate, acetate, lactate, and pyruvate (Figure 5A). Isolated yeast colonies were found on the agar plates supplemented with propionate or butyrate. No yeast colonies or mold was found on the agar plates supplemented with valerate or caproate, which suggested that they have stronger inhibitory effects on microbial growth than other short-chain carboxylates (Figure 5A). Notably, butyrate differentially suppressed the growth of molds and yeasts. Molds did not grow on the agar plates supplemented with butyrate, while the number of yeast colonies reached (9.97 ± 0.45) × 10^5^ CFU/g Daqu, which was much lower than that in the control (Figure 5B).

**Figure 5.**
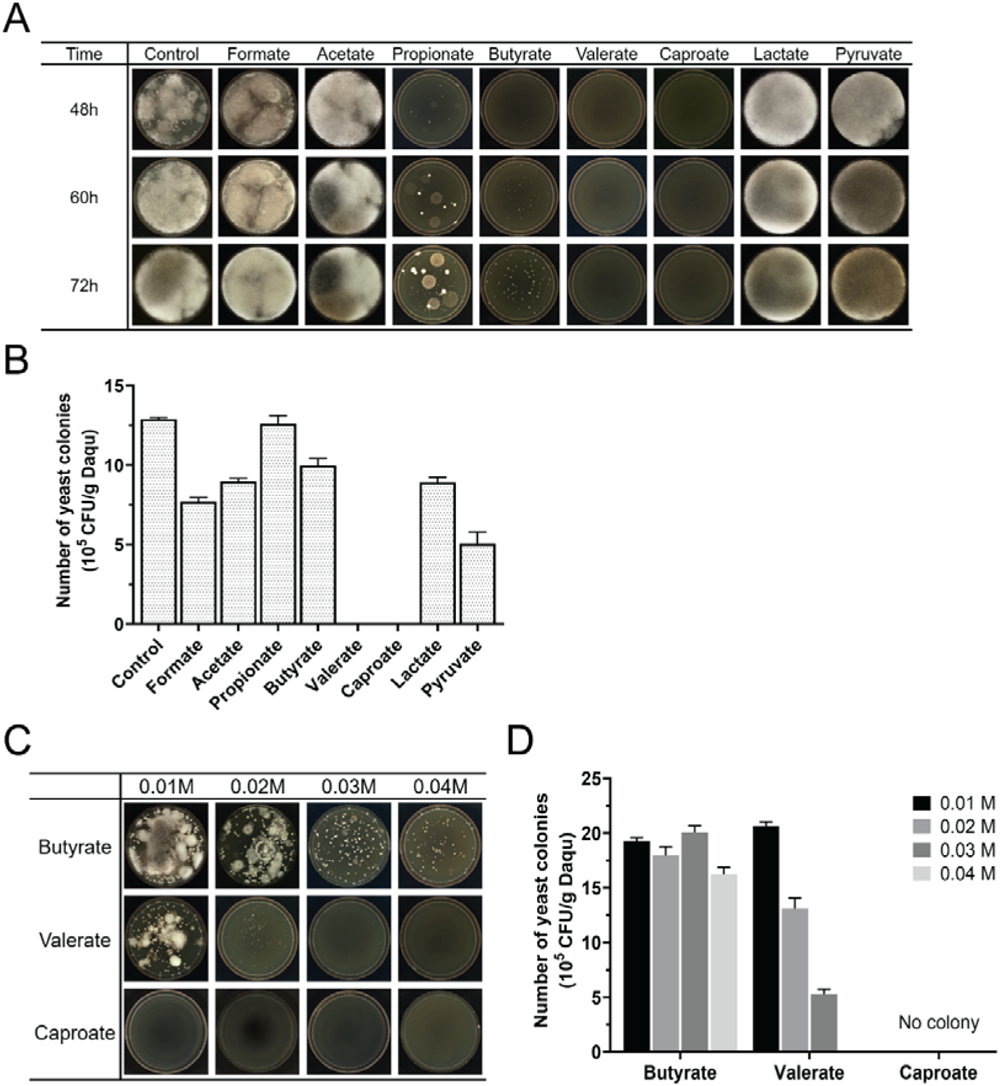
Short-chain carboxylic acids have different inhibitory effects on fungal growth. A, representative agar plates at pH 5.7 and supplemented with formate, acetate, propionate, butyrate, valerate, caproate, lactate, and pyruvate in a final concentration of 0.05 M and control without supplementing short chain carboxylic acid; B, yeast colony counting numbers (CFU) from the agar plates at pH 5.7 and supplemented with various short-chain carboxylic acids and control. C, representative agar plates at pH 5.7 and supplemented with different amounts of butyrate, valerate, and caproate; D, yeast colony counting numbers (CFU) from the agar plates at pH 5.7 and supplemented with different amounts of butyrate, valerate, caproate.

Considering that their inhibitory effect on fungal growth was concentration-dependent, lower amounts of butyrate, valerate, and caproate were added into the MEA medium. For the agar plates that were supplemented with butyrate at a final concentration of 0.03 M, the number of yeast colonies reached (20.07 ± 0.60) × 10^5^ CFU/g Daqu, which is higher than that in the control (Figure 5B, 5C, and 5D). For the agar plates that were supplemented with valerate at a final concentration of 0.01 M, the number of yeast colonies was (20.67 ± 0.35) × 10^5^ CFU/g Daqu (Figure 5B, 5C, and 5D), and decreased at higher concentrations of valerate. Even though lower amounts of caproate were added to the medium, no microbial growth was found on agar plates, indicating the strong toxicity of caproate to the microbes in STD (Figure 5C and 5D).

### 3.6 Acetate, butyrate, and valerate differentially suppress mold and yeast growth in yeast culture media

Based on the results presented above, acetate, butyrate, and valerate were shown to differentially suppress mold and yeast growth and isolated yeast colonies can be counted on the agar plates that were supplemented with these short-chain carboxylates. To validate their inhibitory effects in five different types of yeast medium, acetate, butyrate, and valerate were added at concentrations of 0.05 M, 0.03 M, and 0.02 M, respectively. The pH value was adjusted to 5.0 for the media containing acetate and to 5.7 for the media containing butyrate or valerate. The pH value of the control group was the natural pH of these media without any adjustment. The results showed that all three short-chain carboxylates were effective in inhibiting mold growth on agar plates compared to the control group (Figure 6A). The agar plates supplemented with 0.03M butyrate showed a larger number of yeasts than the control group in all five yeast culture media (Figure 6B), which indicated that it is possible to find a condition that suppresses mold growth and favors yeast growth.

**Figure 6.**
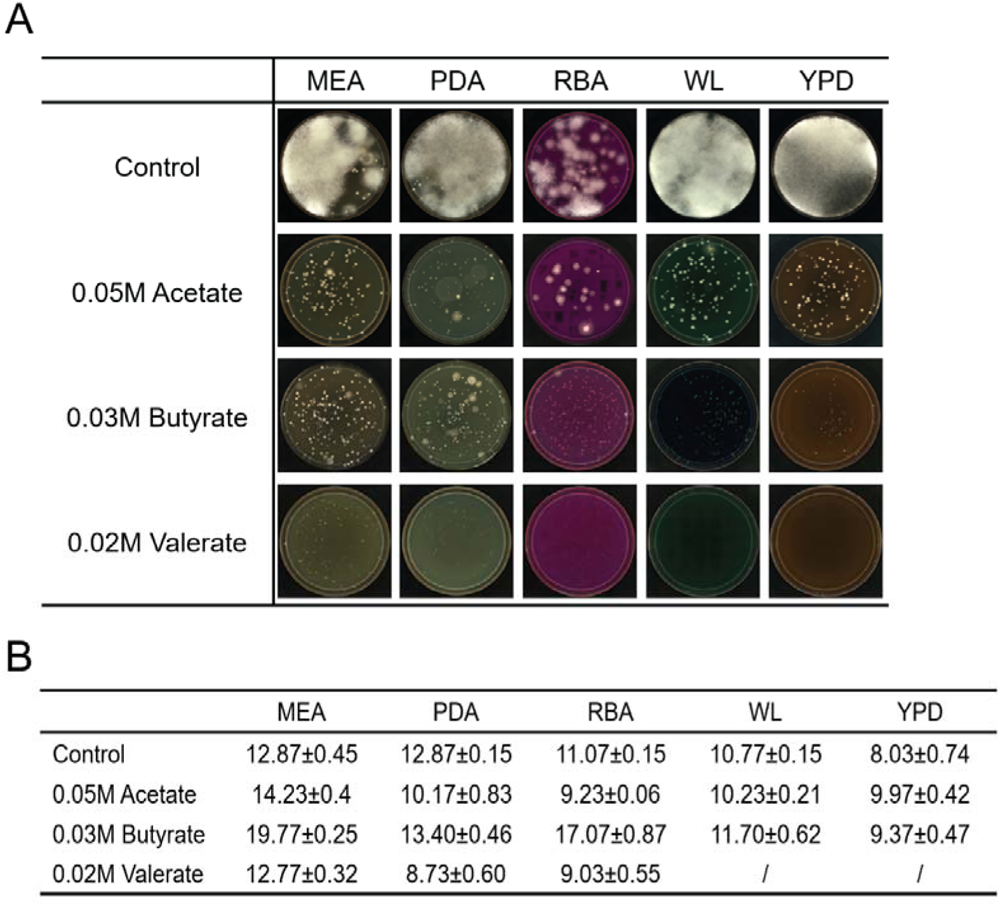
Acetate, butyrate, and valerate differentially suppress mold and yeast growth in yeast culture media. A, representative agar plates supplemented with acetate in a final concentration of 0.05 M at pH 5.0, with 0.03 M butyrate or 0.02 M valerate in a final concentration at pH 5.7; B, yeast colony counting numbers (CFU) from the agar plates supplemented with acetate in a final concentration of 0.05 M at pH 5.0, with 0.03 M butyrate or 0.02 M valerate in a final concentration at pH 5.7.

## 4. Conclusion

In this report, we investigated the effects of short-chain carboxylates on the growth of molds and yeasts in sub-high temperature Daqu. The inhibition of yeast and mold growth on agar plates by short-chain carboxylates depends on both the pH value of the culture medium and the concentration of short-chain carboxylates. Adding certain short-chain carboxylates to yeast culture media differentially regulates the growth of molds and yeasts, which may represent a simple and feasible strategy for improving yeast counting and isolation from mixed culture. It is important to point out that different Daqu starters have different types of microorganisms. An optimization process is needed to find an optimal condition that suppresses mold growth and favors yeast growth.

## Author Contributions

Zhiqiang Ren: Conceptualization, Methodology, Supervision, Funding acquisition, Writing-original draft preparation, Writing-review and editing; Juan Xie: Data Curation, Writing-original draft preparation; Tuoxian Tang, Visualization, Formal analysis, Writing-review and editing; Zhiguo Huang, Supervision, Resources, Funding acquisition. All authors have read and approved the final version of this manuscript.

## Acknowledgments

This study was supported by an open fund project of Liquor Making Biotechnology and Application of Key Laboratory of Sichuan Province (Grant No. NJ2022-07).

